# PanomiR: A systems biology framework for analysis of multi-pathway targeting by miRNAs

**DOI:** 10.1101/2022.07.12.499819

**Authors:** Pourya Naderi Yeganeh, Yue Yang Teo, Dimitra Karagkouni, Yered Pita-Juárez, Sarah L. Morgan, Frank J. Slack, Ioannis S. Vlachos, Winston A. Hide

## Abstract

Charting microRNA (miRNA) regulation across pathways is central to characterizing their role in disease. Yet, current methods reveal only individual miRNA-pathway interactions. We have developed a systems biology approach, *Pathway networks of miRNA Regulation* (PanomiR), that overcomes these limitations to identify miRNA targeting of groups of interacting pathways using gene expression. The approach does not depend on statistically significant enrichment of miRNA target genes in individual pathways or significant differentially expressed genes. Rather, it directly captures differential activity of pathways between states, determining their up-or-down regulation while sensitively detecting biologically-meaningful signals. PanomiR analyzes the co-activity of differentially regulated pathways to determine coordinate functional groups and uses these co-activated grouped pathways to uncover miRNAs that target them. Incorporating both experimentally-supported or predicted miRNA-mRNA interactions, PanomiR robustly identifies miRNAs central to the regulation of disease functions. We applied PanomiR to a liver cancer dataset and showed that it can organize liver cancer pathways and their regulating miRNAs into coordinated transcriptional programs, reflecting the pathogenic mechanisms of hepatocellular carcinoma. PanomiR recapitulated known central miRNAs in liver cancer with a biologically meaningful assignment of pathways under their regulation, unbiased by the number of genes targeted by each miRNA. PanomiR is a granular framework for detecting broad-scale multi-pathway programs under miRNA regulation. It is accessible as an open-source R/Bioconductor package: <https://bioconductor.org/packages/PanomiR>.

## INTRODUCTION

MicroRNAs (miRNAs) are small non-coding RNAs that act as potent regulators of cellular functions and molecular pathways (1). They post-transcriptionally regulate gene expression and can coordinate gene function across distinct pathways. miRNA dysregulation has been shown to be a central component of the pathogenesis of diverse diseases, including neoplastic conditions and Alzheimer’s Disease (2–20). Because miRNAs can target dozens of genes, the characterization of their roles in health and disease requires charting of coordinate co-regulation across heterogeneous molecular cascades and pathways. Despite extensive progress in the field to map the effects of miRNAs on one or more pathway activities (4, 21–27) or the effect of pathway activity on a miRNA, no framework exists for characterization and prioritization of the multi-pathway dynamics of miRNA-orchestrated regulation that form driving transcriptional programs in both healthy and diseased states.

Current best practice for the transcriptomic study of miRNA regulation relies on miRNA-gene or one-to-one miRNA-pathway relationships. miRNA-pathway analysis techniques such as gene set enrichment and correlation are used to detect whether a pathway is potentially regulated by a miRNA (4, 21). Enrichment analyses evaluate the presence (overlap) of targets of a single miRNA in a single pathway, aiming to identify pathways with a higher number of targets than expected by chance (21, 22, 28–32). Alternatively, correlation methods evaluate the association of the expression of a single miRNA with a gene representing the activity of a pathway (4, 33). Table 1 describes some of the most widely used methods for miRNA pathway analyses, their scope and approach. Large-scale functional processes in health and disease coordinate across pathways in multiple ways, including gene-sharing, pathway co-activity, multi-pathway co-regulation, and cross-talk (34–39). Current approaches fail to account for these complex relationships and disease-specific expression dynamics, which in turn limits our ability to detect the potential of a miRNA to regulate highly-specific or broadly-acting gene expression programs.

**Table 1.** Overview of standard miRNA-Pathway analysis methods and PanomiR.

| Method/<br>Reference | Multi-pathway<br>targeting | Pathway<br>activity<br>dynamics | Pathway<br>coordination/<br>interaction | Open-source<br>software | Tissue-specific<br>customization |
| --- | --- | --- | --- | --- | --- |
| **PanomiR**<br>This work | **X** | **X** | **X** | **X** | **X** |
| **miRPath v3**<br>[PMID 25977294] | **X**<br>Meta-analysis |  |  |  |  |
| **Wilk and Braun**<br>[PMID 29294105] |  | **X** |  | **X** | **X** |
| **miRPathDB 2**<br>[PMID 31691816] |  |  |  | **X**<br>precalculated | **X**<br>precalculated |
| **MITHrIL**<br>[PMID 27275538] |  | **X**<br>(via DEG) |  |  | **X** |
| **miRTar**<br>[PMID 21791068] |  |  |  | **X**<br>Web portal non-<br>functional |  |
| **BUFET**<br>[PMID 28874117] |  |  |  | **X** | **X** |
| **miTalos**<br>[PMID 26998997] |  |  |  |  | **X**<br>precalculated |

To uncover how multiple pathways are coordinated by miRNAs to form gene expression programs, we have developed a framework to address existing limitations from a systems perspective. *Pathway networks of miRNA Regulation* (PanomiR) enables discovering central miRNA regulators based on their ability to control coordinate pathways forming a transcriptional program. PanomiR determines if a miRNA concurrently regulates and targets a coordinate group of disease- or function-associated pathways, as opposed to investigating isolated miRNA-pathway events. PanomiR derives these multi-pathway targeting events using predefined pathways, their co-activation, gene expression, and annotated miRNA-mRNA interactions. Its framework (i) captures the activity of pathways and identifies disease-specific differentially regulated pathways using pathway activity profiling, a technique that accounts for overall co-activity of genes and commonly observed biases (40–42); (ii) constructs a co-expression network of differentially regulated pathways (based on a reference of pathway co-expression networks) and deconvolves it into coordinate groups of pathways that act in concert using network clustering algorithms (40–42); (iii) determines miRNAs targeting these coordinate pathway groups using a novel statistical test and pre-determined miRNA-mRNA interactions from experimentally-supported or prediction databases (43, 44). Taken together, these steps produce broad-scale, multi-pathway, and disease-specific miRNA regulatory events (Figure 1).

**Figure 1.**
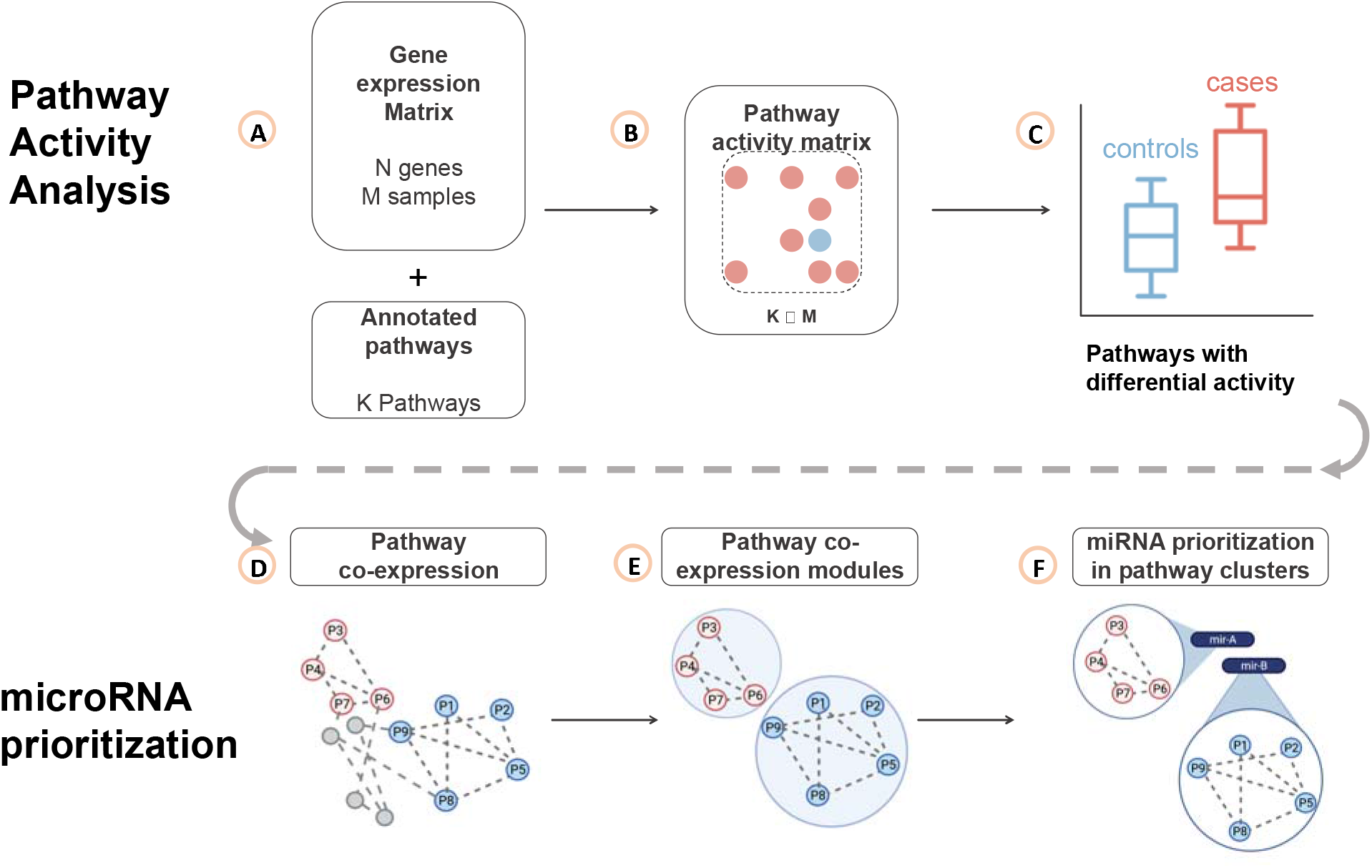
PanomiR workflow. PanomiR prioritizes miRNAs that target coordinate groups of pathways. **(A)** Input gene expression dataset and a set of annotated pathways **(B)** Gene expression data is summarized into pathway activity scores. **(C)** Pathway activity profiles are compared between disease and control subjects to discover differentially regulated pathways. **(D)** Differentially regulated pathways are mapped to the canonical pathway co-expression network (PCxN), where nodes denote pathways and the edges denote correlation of activity scores. **(E)** Within the network of differentially regulated pathways, modules of coordinate pathways are identified using graph clustering algorithms **(F)** miRNAs are prioritized using annotated miRNA-mRNA interactions (known or predicted) for preferential targeting within each cluster of differentially regulated pathways. The outputs of the pipeline are individuals lists of miRNAs with prioritization scores (targeting p-values) per each cluster of pathways.

In order to highlight PanomiR’s ability to detect miRNAs regulating gene expression programs in human disease, we applied it to the hepatocellular carcinoma dataset of The Cancer Genome Atlas (TCGA) comprising 368 primary tumor samples and 49 controls(45). PanomiR unbiasedly detected multi-pathway miRNA regulation events in this well-documented disease and generated a comprehensive framework for evaluating miRNA-associated mechanisms (46). We identified differentially regulated pathways and uncovered their regulating miRNAs in liver cancer, assessed biological relevance of the readouts, and evaluated the statistical robustness of PanomiR using *in silico* experimentation. PanomiR recapitulated known central miRNAs in hepatocellular carcinoma with a biologically meaningful assignment of pathways under their regulation, unbiased by the number of genes targeted by miRNAs. PanomiR is available as an easy-to-use Bioconductor R package, enabling its application in research projects, inclusion into *in silico* tools, and augmentation of analysis pipelines.

## MATERIALS AND METHODS

### Overview and input datasets

The overarching goal of PanomiR is to detect miRNAs regulating multi-pathway condition-associated gene expression programs (Figure 1). PanomiR uses as input a user-provided gene expression dataset (e.g., RNAseq) to quantify pathway activity profiles by utilizing annotated pathway datasets from the Molecular Signatures Database (MSigDB) (Figure 1A-1B)(47). Pathway activity profiles are then compared between two conditions (e.g., cancer *vs* control, wild type *vs* knockout) to identify and prioritize differentially regulated pathways (Figure 1C). To determine broad-scale condition-associated groups of functions, PanomiR constructs a co-activity network of differentially regulated (or disease dysregulated) pathways and deconvolves the network into coherent functional groups using reference pathway co-expression networks; using our previously-described pathway activity methods (Figure 1D-1E) (40–42). Subsequently, PanomiR integrates miRNA-mRNA interactions provided by the user (such as predicted targets from TargetScan (43) or experimentally validated interactions from TarBase (44)) to evaluate miRNA regulatory effects on coordinate pathway groups (Figure 1F). The final output of PanomiR is a ranked list of central miRNAs, together with statistical significance levels for each group of differentially regulated pathways, providing an effective means for identification of pathway groups, and for key miRNA prioritization, ranking, and target detection. PanomiR identifies differentially coordinated transcriptional programs between two conditions to provide a direct prioritization of the miRNAs responsible for their coordination.

### Capturing pathway activity dynamics

Extending the approach developed in our previous methodology, Pathprint (40–42), PanomiR ingests a user-provided gene expression dataset and calculates pathway activity scores to capture pathway functional dynamics (Figure 1B). The scores are proxy values for the activity of genes in individual pathways, which in turn, represent biologically meaningful functional units. By capturing gene expression levels as pathway activity scores, inherent complexity is reduced while tolerance to noise is increased when compared to gene-centric analyses (4, 41, 42, 48). Pathway activity scores leverage the complex inter-relationships and co-activity of genes. They provide the means to examine biological functions in a continuum and detect biological signals where standard differential gene expression analyses fail (4, 33, 40, 42, 48–50).

To capture pathway activity profiles, in a two-step process: (a) we rank genes in each sample in descending order, according to their expression, i.e., the highest expressed gene gets the largest rank-score; (b) we calculate the average squared ranks of genes that belong to a pathway as the activity score. Formally, for a pathway X with n genes, 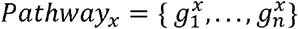 the activity score, *Ac_a,x_*, in sample *a* is:

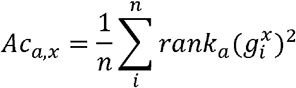

where 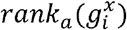 refers to the rank of gene 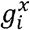 (descending order) in sample *a* based on expression values. We generate activity profiles for each pathway of interest in each sample. Then, pathway profiles are normalized across the input samples (Supplementary Material). PanomiR uses the *canonical pathways* collection from MSigDB as its pathway database reference (47). MSigDB is a carefully curated database that represents non-redundant pathways from established pathway repositories such as KEGG and Reactome (47, 51, 52).

### Detection of differentially regulated disease-associated pathways

PanomiR compares pathway activity profiles between case and control subjects to determine functional dynamics in disease. PanomiR defines differentially regulated pathways by determining statistically significant differences in pathway activity profiles between cases and controls using linear models, implemented in the *Limma* package (Figure 1C) (53). In contrast to enrichment analysis, the linear modeling framework of PanomiR determines the directionality of differential regulation: It defines whether a pathway is upregulated or downregulated in disease subjects (or experimental conditions), and accounts for confounding variables such as batch, sequencing center, or any other fixed effects and continuous covariates. PanomiR outputs an ordered table of differentially regulated pathways along with p-values of differential regulation, adjusted for multiple hypothesis testing using False Discovery Rate (FDR) (54).

### Detection of groups of differentially regulated pathways via their co-expression networks

Dysregulation of an individual pathway is rarely an isolated event since pathways share activity and are often co-regulated. PanomiR accounts for co-regulation to place differentially regulated pathways into groups that represent high-level disease programs by exploiting the *Pathway co-expression network* (PCXN) (42): a reference tool that organizes and assesses the shared activity of pathways (Figure 1D). PanomiR leverages PCxN’s network, generated from a curated dataset of 3,207 expression profiles, providing an independent platform, to query co-activity of all pathways in the MSigDB dataset (42, 55).

PanomiR masks PCxN to contain only the subnetwork of differentially regulated pathways that were identified from the two-group data analysis in the previous step. In the masked network, nodes represent differentially regulated pathways and edges activity-correlation of pathways. PanomiR subsequently identifies densely interconnected differentially regulated pathway subnetworks using graph clustering algorithms (Figure 1E). The default clustering algorithm of PanomiR is Louvain, but PanomiR can use other clustering methods that are available in the igraph R-package (56). The subnetworks denote clusters of highly correlated coordinate groups of differentially regulated pathways driving disease or condition-specific functions.

### miRNA prioritization within clusters of differentially regulated pathways

PanomiR exploits the concept that a coordinate group of disease-associated pathways has common miRNA regulators. Using annotated miRNA-mRNA interactions and an empirical statistical test (Figure 2), it analyzes clusters of differentially regulated pathways, to define central miRNAs, and captures the extent to which the targets of a specific miRNA are present within a group of coordinate pathways. miRNA regulatory events are then identified in three sequential steps (Figure 2): (i) by calculating individual miRNA-pathway overlap scores, (ii) by generalizing miRNA targeting scores to a group of pathways (i.e., a cluster of differentially regulated pathways), and (iii) by estimating the statistical significance of miRNA targeting scores using an empirical approach. The empirical statistical tests are specific to the input dataset, for each miRNA and each cluster of differentially regulated pathways.

**Figure 2.**
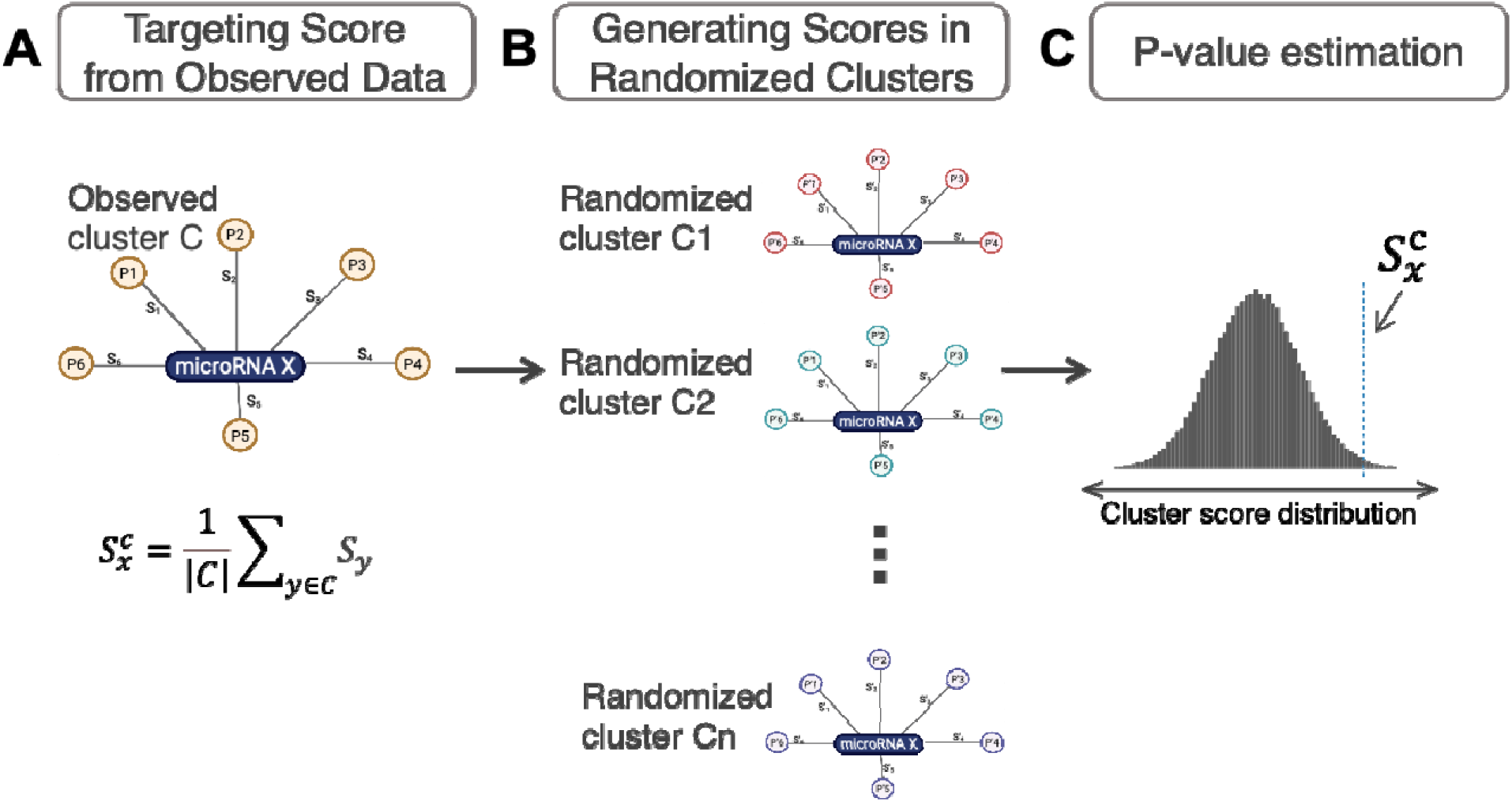
miRNA prioritization from pathway clusters. **(A)** PanomiR generates an observed targeting statistic, *S_X_^C^*, for a miRNA X with respect to *C*, an observed cluster of pathways. The cluster-targeting statistic is an average individual overlap score for each miRNA-pathway pair. Individual overlap scores (e.g., S_1_, S_2_) are functions (inverse normal) of the overlap statistic (Fisher’s exact test) between the miRNA target genes and the pathway member genes **(B)**☐PanomiR generates an empirical distribution of cluster-targeting scores for a miRNA X by randomly selecting a set of pathways and recalculating the cluster targeting score. **(C)** The prioritization p-value is calculated from comparing the observed targeting statistic, *S_X_^C^*, to the null distribution of targeting score for the miRNA X. The p-value is used to rank the miRNAs.

In the first step, PanomiR derives the overlap scores for individual miRNA-pathway pairs using p-values of Fisher’s Exact test, capturing overrepresentation of targets of a specific miRNA in each individual pathway. To make analysis disease-, condition-, tissue-, or cell-type-specific, PanomiR calculates overlap scores using only the genes expressed in the input experiment.

In the second step, an overall targeting score for a given cluster of pathways (Figure 2) is derived. The clusters of pathways are generated in the previous step using PCxN. Formally, for each cluster of differentially regulated pathways, *C*, the targeting score of a miRNA *x* is:

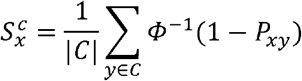

where *Φ*^−1^(.) denotes the inverse of the standard normal cumulative distribution function (CDF) and *P_xy_* denotes the Fisher’s Exact test p-value of overlap between targets of miRNA x and genes of pathway y. The targeting-score, 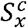, is related to Stouffer’s method (with equal weights) for p-value aggregation. The inverse normal CDF avoids extreme cases in which a miRNA has many targets in one pathway and only a few targets in other pathways in a cluster.

In the third step, the statistical significance of the targeting score 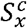 is determined in order to produce cluster-specific lists of miRNAs ranked by targeting p-values. The targeting-score does not constitute, by itself, an unbiased measure of miRNA-targeting as it might depend on the number of targets of a miRNA. To create an unbiased measure, PanomiR also derives an empirical targeting p-value, 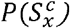, for a score of 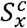. This p-value denotes the probability of observing a larger targeting score from a random cluster of pathways (with |*C*| members) than the one observed. This empirical probability is derived using a bootstrap sampling approach by selecting randomized groups of pathways and re-calculating their cluster targeting score. This approach directly tackles known or unknown biases in gene annotations for miRNA targets, as have been discussed by our group (21) and others (57, 58). The output p-values are then adjusted for multiple hypothesis comparison using the Benjamini-Hochberg False Discovery Rate (FDR) (54).

Given the computational cost of bootstrap sampling, especially to calculate small p-values, PanomiR employs a Gaussian approximation approach to estimate 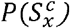. In clusters of large-enough size (>30 pathways), 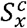 values follow a normal distribution according to the Central Limit Theorem. PanomiR uses pre-calculated Gaussian distribution estimates from 1,000 random *S_x_* values to overcome the computational costs in these cases. In the last step, miRNAs are prioritized based on p-values for targeting each cluster. We provide detailed assessments of the Gaussian estimation and robustness of 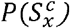-values using a jackknife estimation as Supplementary Materials (Supplementary Figures S3 and S4).

## RESULTS

PanomiR’s utility is presented in a case-study showing its ability to provide a systematic, unbiased, and biologically meaningful determination of regulatory miRNAs. We applied PanomiR to a liver cancer gene expression dataset from TCGA (59). Figures 3 and 4 portray PanomiR’s recapitulation of liver cancer-associated pathways (Table 2), their coordination, and the miRNAs that target them. We found three clusters of differentially regulated pathways in liver cancer representing coherent function of high-level cancer mechanisms: transcription, cell replication, and signaling. PanomiR detected miRNAs that targeted each cluster using either experimentally supported (TarBase v8.0) or predicted (TargetScan v7.2) miRNA-miRNA interactions (Tables 3 and 4). By comparing PanomiR’s results with the relevant literature and with enrichment analysis, we show that PanomiR provides informative and novel biological inference of multi-pathway targeting by miRNAs.

**Table 2.**
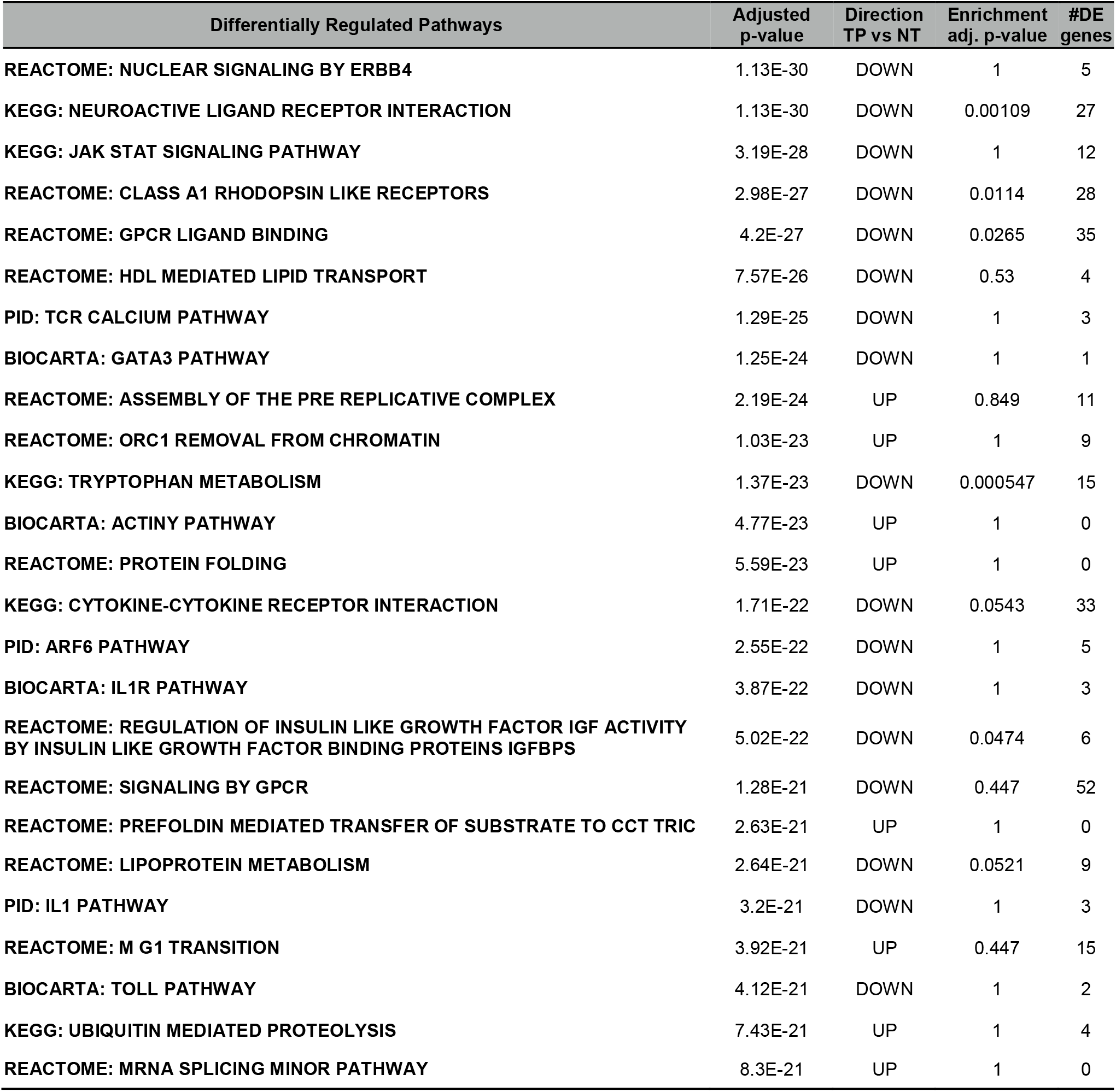
Detection of differentially regulated pathways in liver cancer. Most significant differentially regulated pathways identified by PanomiR according to p-value of differential activity between tumor (TP) and normal tissues (NT). Differential regulation p-values were derived using linear models using the *limma* package by comparing pathway activity profiles of TP vs NT (53). Differential Activation adjusted P-values are for multiple hypothesis testing using FDR. Direction denotes upregulation or downregulation of pathway activity in TP vs NT. Enrichment adjusted p-values for pathways are provided for comparison. Enrichment p-values were derived from the differentially expressed genes (|FC| >1, FDR <0.05). The column “#DE genes” shows the number of differentially expressed genes (TP vs NT) that are present in the pathway.

**Table 3.** PanomiR prioritizes regulatory miRNAs in liver cancer using experimentally-validated interactions. Prioritized miRNAs for each identified pathway cluster, ranked by PanomiR targeting p-value (Figure 2). miRNAs are prioritized based on experimentally validated miRNA-mRNA interaction from TarBase V8.0 (44). Enrichment analysis results are provided for comparison. The column “#Pathways enriched” denotes the number of pathways in the cluster with significant (FDR < 0.25) enrichment in the targets of each miRNA, derived using Fisher’s Exact test.

| Cluster A (n = 65) |  |  | Cluster B (n = 60) |  |  | Cluster C (n = 58) |  |  |
| --- | --- | --- | --- | --- | --- | --- | --- | --- |
| miRNA | # Pathways Enriched | PanomiR Adjusted p-value | miRNA | # Pathways Enriched | PanomiR Adjusted p-value | miRNA | # Pathways Enriched | PanomiR Adjusted p-value |
| hsa-miR-525-3p | 1 | 2.7E-43 | hsa-miR-107 | 40 | 1.92E-22 | hsa-miR-410-3p | 1 | 2.25E-07 |
| hsa-miR-1307-5p | 0 | 1.4E-27 | hsa-miR-124-3p | 38 | 2.04E-22 | hsa-miR-552-3p | 4 | 0.00057 |
| hsa-miR-302c-5p | 0 | 6.22E-24 | hsa-miR-103a-3p | 40 | 3.24E-21 | hsa-miR-5187-5p | 1 | 0.00057 |
| hsa-miR-631 | 5 | 6.61E-24 | hsa-miR-129-2-3p | 37 | 2.31E-17 | hsa-miR-612 | 2 | 0.00254 |
| hsa-miR-663a | 6 | 1.64E-23 | hsa-miR-1-3p | 36 | 2.43E-17 | hsa-miR-198 | 7 | 0.0142 |
| hsa-miR-595 | 4 | 4.03E-23 | hsa-miR-23a-5p | 3 | 8.12E-16 | hsa-miR-621 | 2 | 0.0172 |
| hsa-miR-933 | 1 | 1.04E-22 | hsa-miR-663a | 5 | 3.29E-15 | hsa-miR-199b-5p | 4 | 0.0209 |
| hsa-miR-510-5p | 1 | 1.04E-22 | hsa-miR-449b-5p | 33 | 2.26E-14 | hsa-miR-204-3p | 1 | 0.0249 |
| hsa-miR-5009-5p | 7 | 2.23E-22 | hsa-miR-147a | 41 | 2.26E-14 | hsa-miR-4733-5p | 1 | 0.0503 |
| hsa-miR-2682-5p | 0 | 4.65E-22 | hsa-miR-193b-3p | 35 | 3.26E-14 | hsa-miR-506-3p | 2 | 0.0503 |

**Table 4.** PanomiR prioritizes regulatory miRNAs in liver cancer using predicted interactions. Prioritized miRNAs for each identified pathway cluster, ranked by PanomiR targeting p-value (Figure 2). miRNAs are prioritized based on predicted miRNA:mRNA interaction from TargetScan V7.2 (43). The column “Pathways enriched” denotes the number of pathways in the cluster with significant (FDR < 0.25) enrichment in the targets of each miRNA, derived using Fisher’s Exact test.

| Cluster A |  |  | Cluster B |  |  | Cluster C |  |  |
| --- | --- | --- | --- | --- | --- | --- | --- | --- |
| miRNA | Pathways Enriched | PanomiR Adjusted p-value | miRNA | Pathways Enriched | PanomiR Adjusted p-value | miRNA | Pathways Enriched | PanomiR Adjusted p-value |
| hsa-miR-371a-5p | 0 | 1.06E-38 | hsa-miR-191-5p | 2 | 2.53E-19 | hsa-miR-219a-2-3p | 3 | 1.68E-09 |
| hsa-miR-505-3p.2 | 0 | 8.96E-34 | hsa-miR-892c-3p/hsa-miR-452-5p | 1 | 8.81E-14 | hsa-miR-376c-3p | 4 | 9.51E-09 |
| hsa-miR-1298-5p | 0 | 1.88E-31 | hsa-miR-4424 | 4 | 4.26E-12 | hsa-miR-1249-3p | 0 | 4E-08 |
| hsa-miR-556-5p | 0 | 3.4E-28 | hsa-miR-339-5p | 3 | 8.51E-11 | hsa-miR-3605-3p | 3 | 1.2E-07 |
| hsa-miR-325-3p | 3 | 5.95E-25 | hsa-miR-944 | 0 | 8.51E-11 | hsa-miR-143-3p | 3 | 2.06E-07 |
| hsa-miR-495-3p | 0 | 1.05E-23 | hsa-miR-345-5p | 1 | 2.03E-10 | hsa-miR-514b-5p/hsa-miR-513c-5p | 1 | 6.57E-07 |
| hsa-miR-1278 | 1 | 1.05E-23 | hsa-miR-518c-3p | 0 | 8.89E-10 | hsa-miR-187-3p | 3 | 1.39E-06 |
| hsa-miR-651-5p | 0 | 1.46E-22 | hsa-miR-154-5p | 2 | 3.17E-09 | hsa-miR-625-3p | 1 | 3.06E-06 |
| hsa-miR-323b-3p | 0 | 6.39E-18 | hsa-miR-1251-5p | 1 | 3.47E-09 | hsa-miR-1306-5p | 2 | 5.83E-06 |
| hsa-miR-421 | 0 | 1.16E-17 | hsa-miR-599 | 1 | 3.47E-09 | hsa-miR-873-5p.1 | 3 | 7.82E-06 |

**Figure 3.**
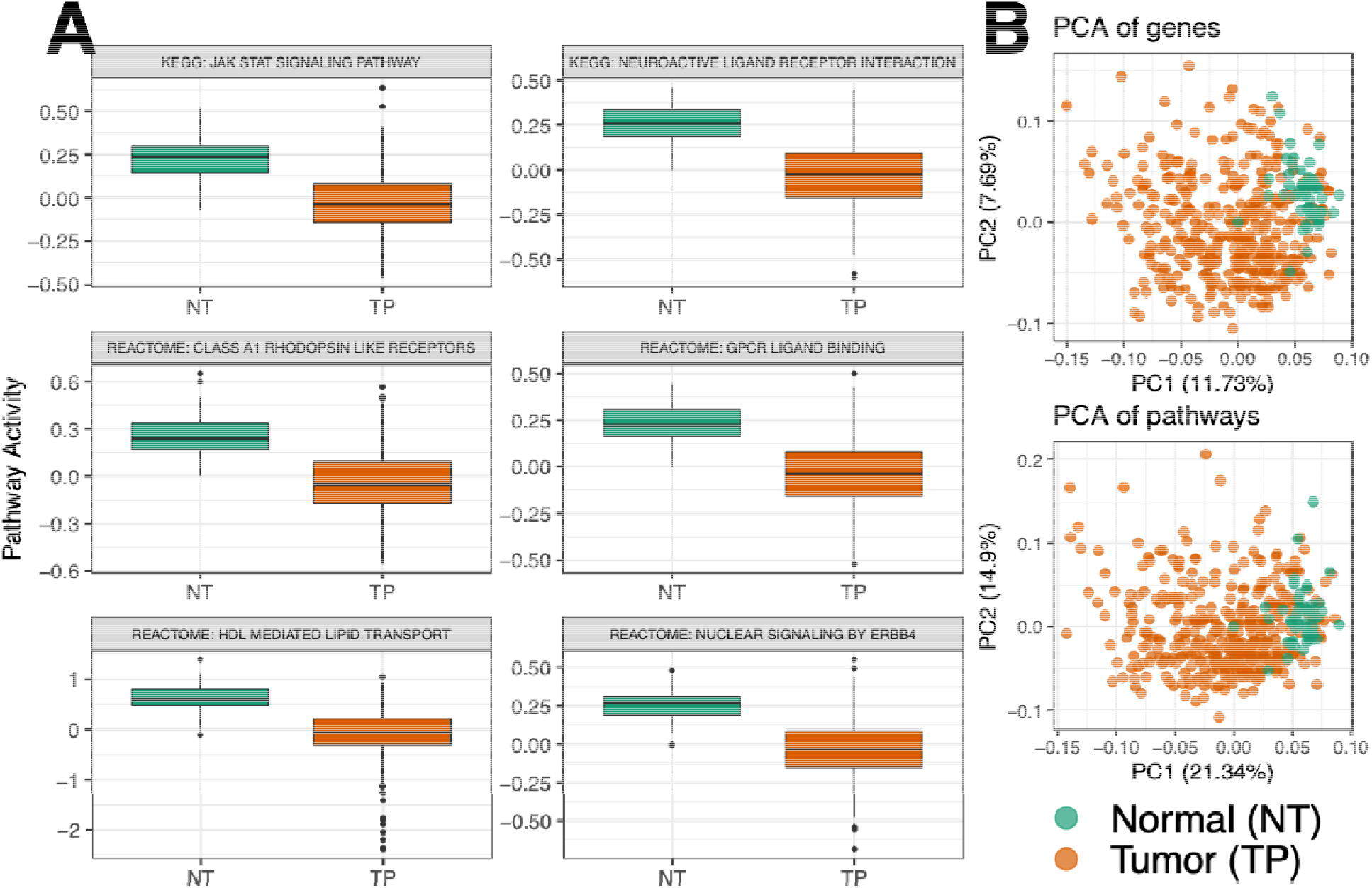
Pathway activity analysis of the Liver Hepatocellular Carcinoma dataset from the Cancer Genome Atlas. **(A)** Detection of differentially regulated liver cancer pathways by comparison of pathway activity profiles between normal tissues (NT) and Tumor primary (TP) samples (45). Boxplots show the most significant differentially regulated pathways selected based on p-values of difference between NT and TP (Table 2). **(B)** Principal component analysis (PCA) projection of the samples based on either genes or pathways. Pathway summarization in PanomiR allows to analyze the activity of pathways in a continuum. PCA of pathways conserves sample groups and captures a higher variation compared to the PCA of genes.

**Figure 4.**
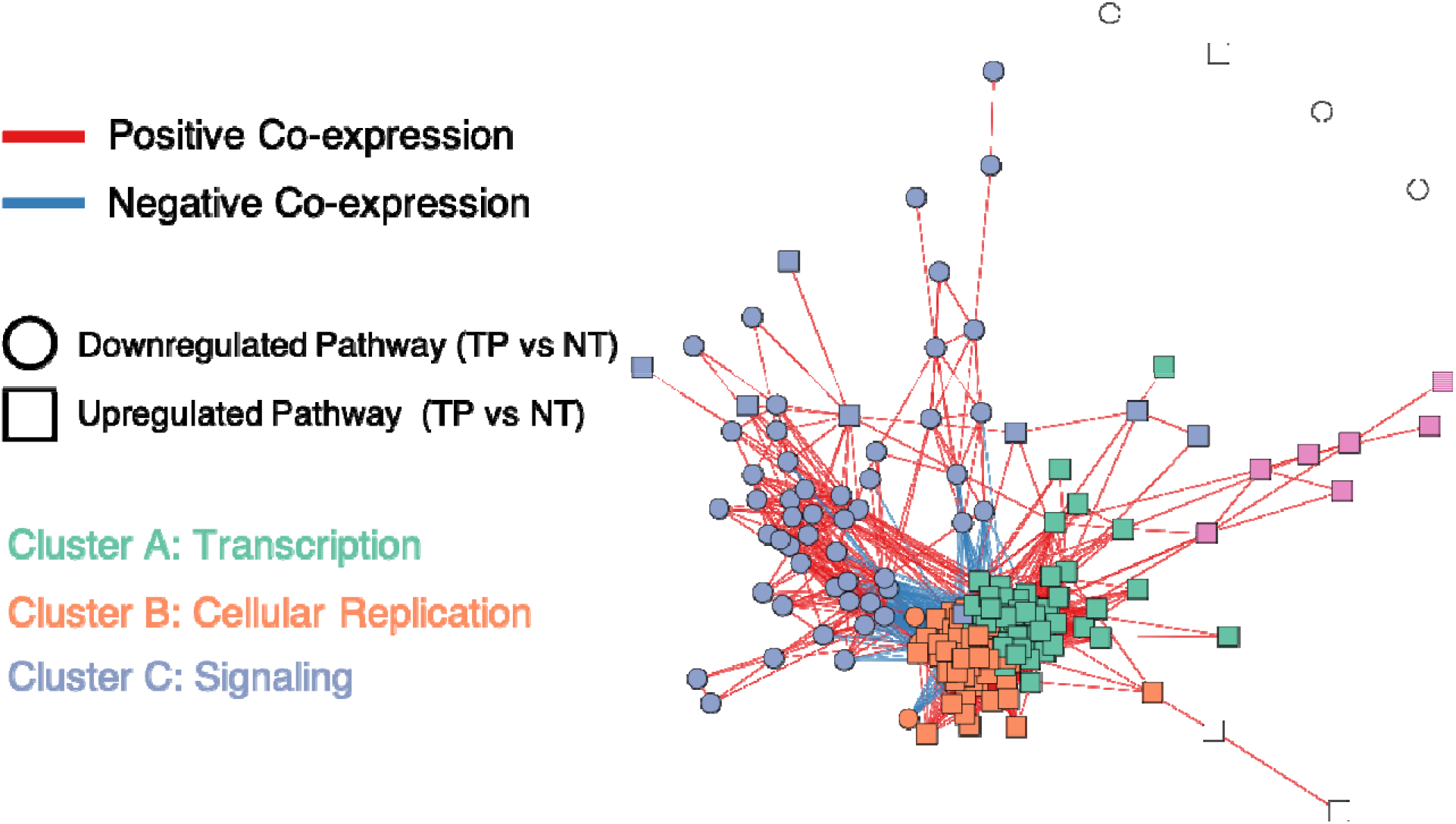
PanomiR deconvolutes coordinate clusters of differentially regulated pathways in liver cancer. The network displays a pathway co-expression map of liver cancer pathways. PanomiR detected three major groups of pathways, defined by direction of differential regulation and clusters of co-expression. The three classes are (i) activation of transcription in tumors (Cluster A) (ii) activation of cellular replication (Cluster B) (iii) deactivation of specific signaling pathways (Cluster C). Each node in the network represents a differentially regulated pathway (Table 2). Edges represent canonical co-expression between two pathways, obtained from an independent compendium of gene expression data, as described in the PCxN method (42). Node colors represent unsupervised network clusters found by Louvain algorithm (83). Clusters were manually labeled according to the functional consensus of their pathways.

### PanomiR detects multiple liver cancer-associated pathways

We generated and compared pathway activity profiles from normal tissues (TCGA Abbreviation: NT, n = 49) and primary solid tumors (TCGA Abbreviation: TP, n = 368) from liver cancer gene expression RNAseq data and using the MSigDB pathway database. PanomiR detected 428 upregulated and 397 downregulated pathways in TP compared to NT (FDR < 0.01, Total pathways 1220; Table 2 and Supplementary Table S1). The large-scale differences in pathway activity profiles closely mirror the differential expression results at the gene level: more than 50% of the genes were differentially expressed based on a similar statistical design (FDR <0.01; n = 7801; total genes= 14212).

Differentially regulated pathways reflected well-established dysregulated functions in liver cancer (Table 2). For example, *NUCLEAR SIGNALING BY ERBB4* was downregulated in TP and activated in NT and has the highest statistical significance among all pathways (Figure 3a, Table 2). Downregulation of ERBB4 in tumors is in concordance with a well-established body of evidence on the roles of ERBB signaling as a tumor suppressor in liver cancer (60, 61). In addition, we found downregulation of *HDL MEDIATED LIPID TRANSPORT* in tumor tissues, corroborated by several reports on lipid disorders in liver cancer including decreased plasma levels of HDL (62, 63). These results suggest the utility of PanomiR in detecting differentially regulated disease functions through pathway activity analysis.

We compared pathway readouts from PanomiR with pathway enrichment analysis of differentially expressed genes from the same liver cancer dataset. Enrichment analysis using Fisher’s Exact Test and comparable cut-offs identified 51 enriched pathways (FDR <0.01, Supplementary Table S2) from differentially expressed genes (differential gene expression: FDR < 0.05; |LogFC| > 1; supplementary material). Of these enriched pathways, 50 were also determined as differentially regulated by PanomiR. Significant overlap between the results suggests that PanomiR recapitulates the majority of enrichment analysis readouts (Fisher’s Exact Test p-value = 3.5e-08). PanomiR also detected liver cancer pathways that were missed by enrichment analysis. For example, the top liver cancer-associated pathway according to PanomiR, *NUCLEAR SIGNALING BY ERBB4,* was not detected by enrichment analysis (p-value = 1), since overrepresentation analysis prioritizes pathways with more DE genes than expected by chance and misses pathways with significantly differential activity between pathways and controls but not increased DE gene proportions. Table 2 and Supplementary Table S1 show several other instances of pathways that were detected by PanomiR but were not identified in the standard enrichment analysis. These results highlight the ability of PanomiR to detect significant functional dysregulation in disease even in absence of significant differential gene expression (Table 2).

### Synthetic data analysis shows PanomiR captures biologically-meaningful signals

To assess the recapitulation of biological signals by PanomiR, we employed two randomization tests (Supplementary Material). In the first, we asked to what extent PanomiR detected differentially regulated pathways in a random assignment of samples to case and control groups in liver cancer (i.e., biologically meaningless classes). PanomiR found a very small number of differentially regulated pathways (0.054 ± 1.2, Mean ± SD) via randomized case/control sample assignment (Supplementary Table S3). In the second test, we interrogated whether the use of biologically meaningful pathways (as annotated in the MSigDB) held advantages versus using randomly assigned gene sets. We generated randomized pathways by permuting gene labels to conserve the overlap structure of the original MSigDB dataset. We found that annotated gene-sets generate a significantly larger number of differentially regulated pathways (one-sided Z test p-value < 3.34 *10^−5^; mean = 693.785 pathways at an FDR < 0.01; sd = 39.1). We also compared the distribution of adjusted p-values from differentially regulated pathways from MSigDB and with that of randomized pathway collections, irrespective of FDR cut-offs. This experiment showed a significant difference between the two scenarios according to a one-sided Kolmogorov-Smirnov test (p-value < 2.86E-18 Supplementary Table S3, Supplementary Figure S1). Biologically meaningful gene sets were more likely to be differentially regulated than randomized pathways and were more likely to capture biological signals.

### Identification of coordinate clusters of differentially regulated pathways

Pathways coordinate and co-regulate through various mechanisms, including gene sharing. To detect coordinate groups of differentially regulated pathways, we used the Pathway Co-expression Network (PCxN), where edges represent precalculated correlations between pathways based on independent gene expression data (42). We mapped the 200 most statistically significant differentially regulated pathways onto the PCxN network and performed Louvain clustering to identify coordinate pathway groups. PanomiR identified 3 major clusters of differentially regulated pathways with consistent functions (Figure 4).

The largest cluster of differentially regulated pathways (Cluster A) contained pathways upregulated in TP such as *SPLICEOSOME, PROTEASOME, TRANSLATION, RNA POLL II TRANSCRIPTION, and SIGNALLING BY WNT* (Supplementary Table S4). Wnt signaling activation is a critical mechanism for transformation of precancerous lesions into liver cancer through proliferation (64). The second largest cluster (Cluster B) contained terms related to cell cycle and proliferation (Figure 4, Supplementary Table S4). The third cluster (Cluster C) contained liver cancer-associated signaling pathways that were either down or upregulated in TP vs NT with terms related to ERBB signaling, IL signaling, and NOTCH signaling (Supplementary Table 4). Differentially regulated pathways within clusters A and B showed a coherent direction of differential regulation in cancer vs normal tissues, suggesting a coordinate multi-pathway dysregulation in driving high-order functions. We validated the robustness of pathway clustering using a variety of parameters and algorithms (Supplementary Figure S2). The results indicate that PanomiR successfully deconvolves distinct groups of differentially regulated pathways that represent higher-order functional programs of liver cancer.

### Detection of regulatory miRNAs that target clusters of differentially regulated pathways

We evaluated whether the coordinate clusters of differentially regulated pathways have common miRNA regulators. In our case study, we examined separately experimentally supported (TarBase v8.0; >500K interactions) and predicted miRNA-mRNA interactions (Targetscan v7.2; >113K interactions) to detect miRNAs that target each cluster of differentially regulated pathways (Tables 3 and 4, Supplementary Tables S5 and S6). Our results showed that PanomiR identified distinct miRNAs for each cluster of liver cancer-associated pathways.

With the use of experimentally supported interactions, PanomiR detected 202, 104, and 1 miRNA regulators in clusters A, B, and C respectively (FDR < 10^−5^, Table 3, Supplementary Table S5). These included known liver cancer-associated miRNAs with consistent modes of action with their targeted pathway clusters. Cluster A was targeted by miR-525-3p, miR-1307, miR-631, and miR-663a– these miRNAs have been previously shown to have a role in tumor migration and invasion (65–68). Cluster B was targeted by miRNAs with established roles in regulating cell-cycle in liver cancer including miR-107, miR-124-3p, and miR-103a-3p. For example, miR-107 is a P53-associated regulator of cell cycle and proliferation, elevated in early-stage liver cancer (69–72); miR-124-3p is a tumor suppressor that regulates proliferation and invasion in liver cancer by inducing G1-phase cell-cycle arrest (73, 74); miR-103a-3p is a promoter of proliferation that is highly dysregulated in liver cancer (75). In cluster C, we found miR-410-3p as a central regulator of the relevant module. This miRNA has been shown to be a circulating biomarker of distant metastasis into the lung and the liver (76, 77), it also regulates adenomas via signaling pathways such as MAPK, PTEN/AKT, and STAT (78, 79). In cluster C, we also found a significant targeting role for miR-552-3p, which has been associated with liver cancer and regulates various hallmarks of cancer (80). Supplementary material provides an examination of the relationship between PanomiR miRNAs with DE miRNAs in TP vs NT. While we did not find a significant association between prioritization by PanomiR and differential expression, PanomiR attributes distinct DE miRNAs to distinct groups of pathway-targeting events– providing a knowledge-driven approach for functional characterization of data-driven disease miRNAs. Our results establish that PanomiR successfully detects key regulating liver-cancer miRNAs and their downstream differentially regulated pathways.

PanomiR was also assessed using predicted miRNA-mRNA interactions (43). Although PanomiR detected multiple liver cancer-associated miRNAs from predicted interactions, the set of prioritized miRNAs were different than that of experimentally supported interactions (Table 4, Supplementary Table S6). For example, PanomiR prioritized miR-299-3p in cluster C, a regulator of IL and STAT signaling pathways in liver cells, which have several associated annotated pathways in cluster C (81). Supplementary tables and results provide information on processing predicted interactions and additional evaluations of PanomiR using varying parameters for selection of predicted interactions. Our results suggest that predicted and experimentally validated miRNA interactions databases produce complementary results, and both should be considered for the downstream analysis of transcriptomic data.

We compared PanomiR’s results with a standard miRNA-pathway enrichment analysis in our case study (Tables 3 and 4). For comparative purposes, we employed three extensions of miRNA-enrichment analysis tests to adapt to multi-pathway scenarios. (a) We initially extended enrichment analysis to a group (cluster) of pathways by interrogating the number of pathways within a given cluster that were significantly enriched for targets of a miRNA. For example, if the targets of a miRNA, x, are significantly enriched in five pathways within a group of pathways, the miRNA gets a targeting score of 5. (b) We used Stouffer’s method to obtain one single p-value that combines enrichment p-values of a miRNA within all pathways in a cluster. (c) We used Fisher’s p-value aggregation method to combine all enrichment p-values of a miRNA as an alternative of Stouffer’s method.

PanomiR successfully detected liver cancer-associated miRNAs that were not prioritized by extended enrichment tests. When using experimentally supported miRNA-mRNA interactions, enrichment analysis of cluster A revealed miR-525-3p as enriched in only 1 and miR-1307-5p in none out of 65 pathways (Table 3). When using predicted miRNA-mRNA interactions, PanomiR detected several miRNAs that were not detected by the extended enrichment analysis (Table 4). It is of note that the enrichment tests (Tables 3 and 4) used a highly-relaxed FDR threshold (FDR <0.25) to enable a more sensitive detection. Using a conservative FDR cut-off (e.g., FDR < 0.05) would have retained an even lower detection rate of miRNAs. The results suggest that (a) PanomiR can detect liver cancer-associated miRNAs that are not detectable by simple enrichment tests, and (b) a subset of critical liver cancer miRNAs can be detected only by analyzing a group of pathways, and not by examining individual pathways.

Enrichment analyses are biased towards detecting miRNAs with a larger number of targets (80). We examined the relationship of the number of targets of a miRNA with its prioritization ranking by PanomiR or enrichment analysis extensions (Table 4). The enrichment ranking of miRNAs significantly correlated with their number of gene targets while PanomiR was unbiased to this number. Figure 5 displays the bias of enrichment analysis (including Stouffer’s and Fisher’s extensions) towards prioritizing miRNAs with more targets in cluster A, while miRNAs with a small number of targets did not rank highly. PanomiR did not show a correlation between ranking according to PanomiR and the number of targets (Figure 5), suggesting its ability to prioritize miRNAs irrespective of the number of their gene targets. Additional evaluation of unbiased and robust miRNA prioritization by PanomiR is provided in the Supplementary material. Using jackknifing and bootstrapping, we showed that PanomiR miRNA prioritization is rather based on collective targeting of all pathways and is not driven by individual pathways

**Figure 5.**
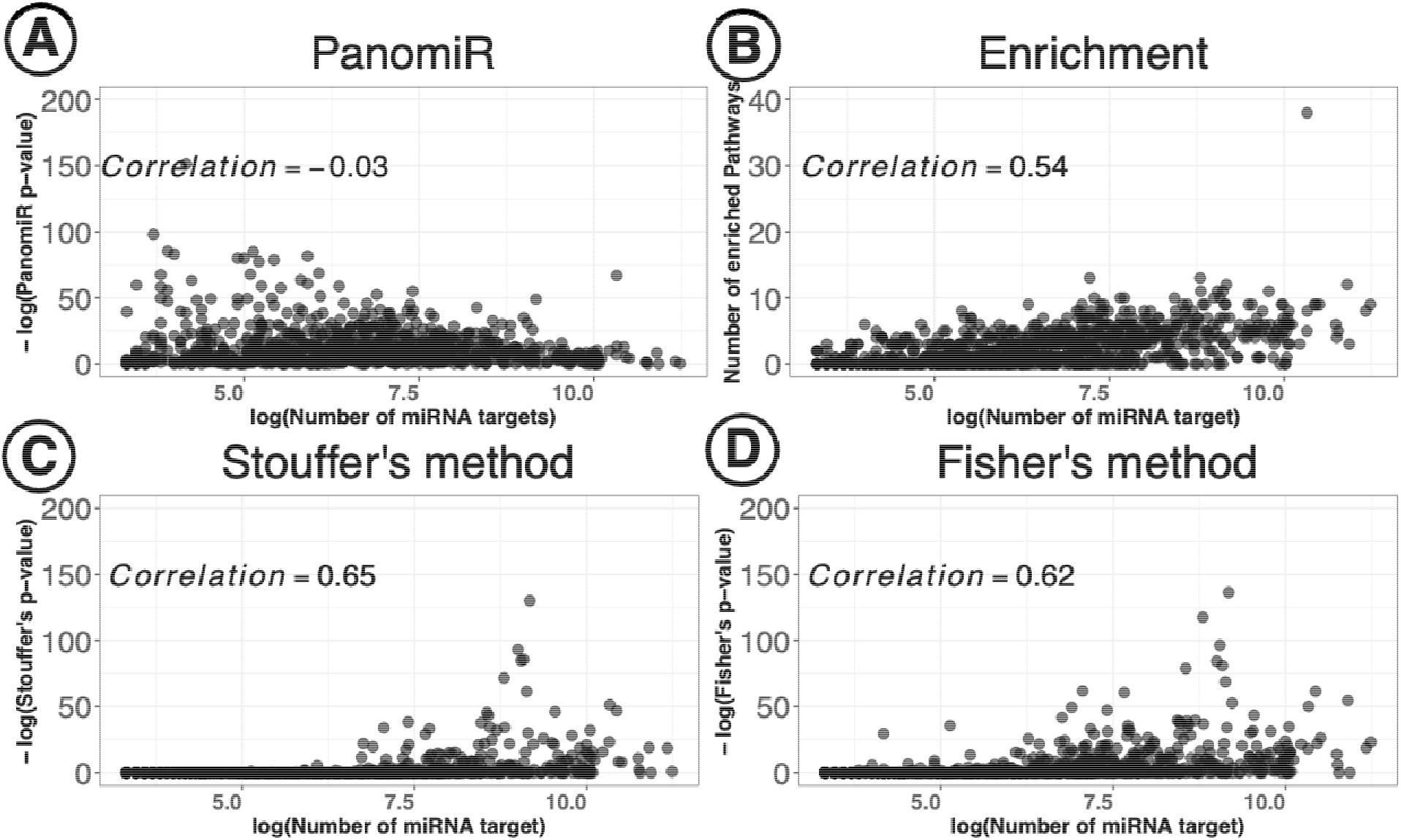
Unbiased prioritization of miRNAs by PanomiR. PanomiR prioritizes miRNAs with either a small or large number of annotated targets. In contrast, enrichment-based miRNA-prioritization methods are biased towards prioritization of miRNAs with larger numbers of gene targets. The figure displays correlation analysis of miRNA-prioritization rankings with the number of gene targets in Cluster A of the liver cancer dataset. Each point represents a miRNA annotated in the TarBase dataset. **(A)** Spearman correlation analysis did not find a significant association between the number of targets and the prioritization ranking of miRNAs by PanomiR (correlation −0.03). **(B)** The number of enriched pathways for a miRNA significantly correlated with its number of gene targets. We also observed a significant correlation between the number of a miRNA’s targets and its prioritization ranking based on **(C)** Stouffer’s method and **(D)** Fisher’s method for aggregation of enrichment p-value. X-axis denotes the log number of gene targets of miRNAs based on experimentally-validated miRNA-mRNA interactions from the TarBase database (44). Y-axes in panel b represents the number of significantly enriched pathways (Adjusted p-value < 0.25, Table 3).

## DISCUSSION

We have built PanomiR, a framework able to determine miRNA regulation of multiple coordinately regulated pathways. Most of the existing tools for miRNA-pathway analysis are focused on one-to-one miRNA-pathway relationships without the ability to infer relationships between miRNAs and groups of co-regulated pathways. Previous studies use of p-value integration methods to address multi-pathway analysis, but none of them determine pathway co-activity/coordination and account for expression dynamics (82). PanomiR addresses these challenges by deconvolving gene expression datasets into coordinate groups of pathways with condition-associated dynamics and by measuring the extent to which miRNAs target these groups. In the case study of the liver cancer dataset, PanomiR captured large-scale features of cancers such as dysregulated transcription, cellular replication, and signaling (Figure 3). These clusters represent coherent higher-order functional units that recapitulate specific, yet central, disease mechanisms.

The use of pathway activity profiles is a key component of PanomiR; It sensitively detects differentially regulated pathways and provides granularity in definition of coordinate functional groups (Figures 2 and 3, Table 2). In a case study, PanomiR detected critical known liver cancer pathways even with few associated differentially expressed genes (Table 2). Pathway activity profiling in PanomiR also facilitated the understanding of the directionality of pathway (de)activation in disease states. Methods that use pathway activity scores are often limited in generating explainable and biologically meaningful pathway activity profiles as they may use nonlinear dimensionality reduction approaches (3). PanomiR fills this gap by providing biologically meaningful measurements of changes in pathway activity profiles where a higher (or lower) pathway activity indicates a higher (or lower) overall activity of associated genes. Pathway activity profiles in PanomiR are directly comparable and translatable across different datasets, which makes it possible to leverage co-expression of pathways to detect disease-specific functional dynamics and themes across datasets, platforms, and species (40).

Deconvolution of coordinate pathway groups allowed PanomiR to detect miRNA regulatory events in liver cancer robustly and unbiasedly, many of which were not detectable by conventional analyses (Figure 4, Tables 3 and 4). PanomiR’s prioritized miRNAs have distinct roles in liver cancer, concordant with the functional characteristic of the pathway clusters that they were discovered from. For example, miR-107 regulates cell cycle and proliferation and targets cluster B which includes cellular replication pathways (Figure 4, Table 3). These results highlight the ability to identify miRNAs that consistently target groups of pathways even with only a few targets from each pathway. Our results demonstrate that PanomiR can robustly detect miRNAs that regulate the broad, yet specific, gene expression programs of liver cancer.

The complete landscape of miRNA-mRNA binding events is currently unknown. This gap contributes to the discrepancy in miRNA prioritization based on the background dataset of miRNA-mRNA interactions (Tables 3 and 4). By using 113K high-confidence predicted miRNA interactions (TargetScan) and more than 500K experimentally supported (TarBase) miRNA targets, PanomiR discovered informative and complementary miRNA regulatory events (Tables 3 and 4). Users have the ability to tailor the background datasets (miRNA-mRNA integration or pathway gene sets) to their study design and research questions. We have made PanomiR flexible to heterogeneous miRNA-mRNA interactions and gene-expression datasets. PanomiR can be expanded (in future development) to co-expression analysis of miRNAs and pathways, which has been proposed to provide informative pointers to biological programs of diseases (3).

In summary, PanomiR is a systems biology framework to study differentially regulated pathways, their co-activity, and their regulating miRNAs. It accounts for co-expression of pathways and disease-specific expression dynamics to identify miRNA-regulatory events, providing an advance over the current practice of studying static and isolated miRNA-pathway interactions. PanomiR is available as an open-source R/Bioconductor package for the use of the community.

## AVAILABILITY

PanomiR is available as a Bioconductor package. The development version of PanomiR can be accessed via <https://github.com/pouryany/PanomiR> and <https://bioconductor.org/packages/PanomiR>. Additional scripts and analyses, specific to this manuscript are available via <https://github.com/pouryany/PanomiR_paper>.

## ACKNOWLEDGEMENTS

We would like to thank Dr. Isabel Castanho and Ms. Saloni Shah for valuable feedback. We thank the Cure Alzheimer’s Foundation and Harvard Medical School for their sponsorship.

