## Supplementary Material for "PanomiR: A systems biology framework for analysis of multi-pathway targeting by miRNAs"

**Supplementary Methods**

**Reagents and test case datasets**

We retrieved a dataset of 417 samples of Liver Hepatocellular Carcinoma (LIHC) patients and controls with concordant miRNA and gene expression profiles from the Cancer Genome Atlas (TCGA) project via the *TCGAbiolinks* R-package (Mounir et al. 2019). We performed downstream analysis including differential expression and enrichment analysis after normalization and preprocessing miRNA and gene data.

**Gene expression processing**

Gene expression profiles from RNA-Seq datasets are processed prior to generating the pathway summary statistic as follows: (a) raw RNA-Seq counts are retrieved and filtered to only include genes with minimum of 1.0 counts per million reads present in at least 20% of the samples. (b) Samples are then quantile-normalized while adjusted for GC-content and gene-length using Conditional Quantile Normalization (CQN) (Hansen, Irizarry, and Wu 2012). GC-content and gene-length are known sources for capturing bias in RNA-Seq studies, and this can affect ranking of genes in RNA-seq experiments (Hansen, Irizarry, and Wu 2012; Benjamini and Speed 2012).

**miRNA expression processing**

miRNA expression profiles for the TCGA liver cancer dataset (Cancer Genome Atlas Research Network. Electronic address: and Cancer Genome Atlas Research Network 2017). were retrieved using the *Firebrowse* portal. miRNA counts were filtered for entries with reads per million (RPM) values of more than 1 in at least 20 percent of the samples. Differential expression was performed on log (RPM + 3) values using a linear model, adjusting for batch, using the *limma* package (Ritchie et al. 2015).

**Characterization of differentially-expressed miRNAs**

A common query for miRNA analysis tools is to functionally characterize differentially expressed miRNAs or miRNAs that target differentially expressed genes. PanomiR performs this task by assigning DE miRNAs to the coordinated clusters of dysregulated pathways. We used PanomiR to characterize DE miRNAs between NT and TP in the liver cancer dataset from TCGA (FDR < 0.01, Supplementary Table S7). For clusters A, B, and C, we PanomiR p-values of DE miRNAs using an FDR < 0.001 criterion.

PanomiR uniquely attributes DE miRNAs to distinct clusters. For example, PanomiR attributes 60 DE-miRNAs to Cluster A using TarBase interactions, including known liver cancer-associated miRNA miR-29b-1-5p, miR-29b-2-5p, miR-34a-3p, miR-122-3p, and miR-221-5p (Callegari et al. 2015; Morishita et al. 2021) (Supplementary Table S8). Similarly, PanomiR uniquely attributes 30 DE-miRNAs to Cluster B including known liver cancer-associated miRNA miR-101-5p, miR-29c-5p, and several miR-26 species (Supplementary Table S8) (Callegari et al. 2015; Morishita et al. 2021). Using experimentally-supported interactions, no miRNAs were attributed in DE to Cluster C. Our results show that PanomiR can attribute DE miRNAs to different groups of pathways with distinct functionalities, providing a higher-order functional characterization.

**Experimentally validated miRNA-mRNA interactions**

miRNA targets were retrieved in July 2019 from the TarBase V8 collection of experimentally validated miRNA:mRNA pairs (Karagkouni et al. 2018). TarBase database was retreived through an access-request application through TarBase portal.The entries in TarBase were limited to positive pairs in *Homo sapiens* species, representing 526,658 interactions containing 1077 unique miRNA species. miRNA:mRNA pairs were limited to those with annotated ENSEMBL ID for genes and the ClusterProfiler package was used to map the ENSEMBL IDs of into ENTREZ ID (Yu et al. 2012).

**Predicted miRNA-mRNA interactions**

miRNA targets were retrieved in July 2021 from the TargetScan V7.2 collection of predicted miRNA:mRNA interaction pairs (Agarwal et al. 2015). Summarized predictions (approximately 23 million human Gene:miRNAfamily pairs) were retrieved. We only used gene:miRNAFamily pairs with negative *context++* scores (Agarwal et al. 2015). The interaction pairs were limited to *broadly conserved, conserved,* and *poorly conserved* categories. miRNA families that belonged to star miRNAs (complementary miRNA sequences) or RNA fragments were excluded from further analysis*.*

We deployed four strategies to generate a list of miRNA targets. In the first strategy we selected all pairs of conserved and broadly conserved miRNA families along with gene targets with at least one conserved 3’UTR binding site. For poorly conserved miRNA families, we selected pairs with a *Cumulative weighted context++ score* of -0.2 or less. This approach is consistent with the outputs of the online portal of the TargetScan.

In other strategies, we filtered miRNA:mRNA pairs based on the *Cumulative weighted context++ score,* agnostic to conservation status of the miRNA family (Paraskevopoulou et al. 2018). These strategies do not compromise the predictive power of the algorithms and do not generate more false-positives when compared to using conserved targets only. The strategies use different context++ score filtering thresholds (Second Strategy: <-0.1, third strategy <-0.2, and fourth strategy <-0.3). Supplementary Table S10 shows the readouts from all four strategies in the liver cancer dataset. We used the fourth strategy (context ++ score <-0.3) for the final analysis.

**Pathway Datasets**

The pathway collection references included 1329 canonical pathways from the Broad Institute’s Molecular Signatures Database (MSigDB, C2, V6.2, downloaded July 2018) (Liberzon et al. 2011). The MSigDB collection consists of pathways from Kyoto Encyclopedia of Genes and Genome (KEGG), BioCarta, Reactome, the Pathway Interaction Database (PID), the Matrisome Project, Signal Transduction Knowledge Environment, and Signaling Gateway (Kanehisa and Goto 2000; Nishimura 2001; Jassal et al. 2020; Schaefer et al. 2009; Naba et al. 2012; Gough 2002; Saunders et al. 2008). For our final analysis, we used pathways with at least 10 expressed genes in the liver cancer dataset.

**Pathway Co-expression Network (PCxN) Processing**

A Pathway Co-expression Network was constructed using MSigDB V6.2 canonical pathways according to the methodology described in (Pita-Juárez et al. 2018). We limited the PCxN to high-confidence edges where the absolute value of the correlation coefficient was larger than 0.1 with an adjusted correlation p-value of < 0.05. The final network was partitioned using the Louvain algorithm into multiple clusters. PanomiR uses a flexible implementation that allows for use of different clustering algorithms including Louvain, edge-betweenness, and Infomap (49–51). Clusters were labeled according their sizes, with Cluster 1 being the largest. The largest three clusters were renamed A, B, and C for clarity.

**Assessing DAP-PCxN consistency**

To assess whether clustering of dysregulated pathways via PCxN is dependent on the number of input pathways, we generated DAP-PCxN in multiple settings choosing varying number of input top statistically significant DAP. In each scenario, X number of top differentially activated pathways were selected and mapped to PCxN and clusters were identified using the Louvain algorithm, with X ranging from 150 to 450 pathways, in 50 pathway increments. We compared correspondence at each setting using the Jaccard index for the top three clusters (Supplementary Figure S2). Our results show that clusters are consistent across a range of choices for the number of input pathways. Hierarchical clustering using the Jaccard index similarity matrix showed one-to-one correspondence between clusters at different settings.

**Normalization of Pathway Summary Statistics**

The pathway profiles, ${Ac}_{a,x}$, are normalized in each sample using the z-transformation, $Z_{a,x}= (Ac_{a,x} - \mu_{a})/\sigma_{a}$, where $\mu_{a}$ and $\sigma_{a}$ denote the mean and the standard deviation of all pathway summaries in sample “*a*”. To ensure that the summary statistics were not influenced by changes in a single gene and to increase robustness, we used a trimmed mean calculation for $Ac_{a,x}$ by removing the top 2.5% highest and 2.5% lowest expressed genes from the pathway.

**miRNA-pathway association scores**

To calculate aggregated scores, $S_{x}^{c},$ PanomiR uses Fisher’s Exact test to generate individual scores. For every miRNA $x$ and pathway $y$, the test will generate an enrichment p-value, $P_{xy}$, which represents the probability of observing a larger overlap between miRNA $x$ ‘s gene targets and genes in pathway $y$. To facilitate computation, whenever a miRNA $x$ does not have target in a pathway $y$, the enrichment p-value $P_{xy}$ is substituted with the largest non-1 enrichment p-value in all miRNAs and all pathways. We performed enrichment analysis only on genes that were expressed in the liver cancer dataset to create a tissue-specific analysis.

**Pathway enrichment analysis of gene expression data**

Differentially expressed (DE) genes between tumor and normal tissues were determined using the normalized gene expression data. Batch effects were adjusted using linear models in the limma package. DE genes were selected using an absolute-value-of-fold-change > 1 cut-off and an FDR< 0.05. Enrichment analysis was performed using clusterProfiler (Yu et al. 2012). Enrichment analysis of MSigDB canonical pathways was performed using the same background of pathways with which the activity profiles where generated.

**Significance of aggregated p-values and miRNA prioritization**

Bootstrap sampling was used to determine the statistical significance of the aggregated scores, $S_{x}^{c},$ against randomized pathway clusters of the same size. If we consider a randomly derived pathway set, $C^{'},$ from the set of all pathways Y, the bootstrapped p-value, $P(S_{x}^{c})$, for miRNA $x$ can be found using a similar formula

$$P\left( S_{x}^{c} \right)=P\left\{ C^{'}\in Y \right| S_{x}^{c^{'}}> S_{x}^{c}\}$$

We can then estimate $P(S_{x}^{c})$ by generating different $C^{'}$and evaluating the number of scores larger than the observed one. Given that this schema can be computationally expensive to estimate small p-values, we use an approximation from a Gaussian distribution to estimate $P\left( S_{x}^{c} \right)$*.* We generate 1000 random clusters of pathways ($C^{'}$) and estimate the mean and standard deviation of the aggregated association scores ($S_{x}^{c^{'}}$). Note that $S_{x}^{c^{'}}$ values estimate the mean of $\Phi^{-1}(1-P_{xy})$ for all pathways. According to central limit theorem, given large enough (|C|), the aggregated association scores follow a normal distribution and we use this property to estimate $P(S_{x}^{c})$.

**Assessment of estimated Gaussian distribution**

By bootstrapping 10000 of these random aggregated p-values in our test case dataset, we calculated the probabilities of significance, $P(S_{x}^{c})$, for each miRNA $x$. We compared bootstrapped p-values with Gaussian estimated p-values from 1000 permutations. We evaluated concordance using correlation analysis.

**Assessment of robustness of miRNA prioritization p-values with jack-knifing**

We examined whether $P(S_{x}^{c})$ were dependent on any single pathway in the cluster. To determine the robustness we used jack-knifing to remove a pathway at a time from the cluster and recalculated the probability of significance. We obtained the mean of all the recalculated probabilities and termed this mean as the jack-knifed probability of significance. For cluster $C$ and miRNA $x$, the jack-knifed probability of significance, ${JP}_{x}^{c}$, can be represented with the formula

${JP}_{x}^{c}=\frac{1}{|C|}\sum_{y\in C} \frac{1}{|C|}[\sum_{y^{'}\neq y, y^{'}\in C} \Phi^{-1}(1-P_{xy'})]$

We compared the jack-knifed estimate to those obtained from the whole cluster using a correlation analysis to evaluate robustness. The comparisons were generated based on the p-values of the test case dataset.

**Stability of PanomiR miRNA prioritization**

We hypothesized that PanomiR robustly prioritizes miRNAs from a group of pathways as a whole, and its prioritization is not driven by single pathways. We used a jackknifing schema to validate the robustness of PanomiR by recalculating PanomiR p-values while repeatedly excluding pathways from clusters one at a time. We calculated a jackknifing p-value by averaging the outputs of PanomiR in all repetitions. We hypothesized that if the PanomiR p-values depended on single miRNA-pathway relationships, then the jackknife p-values would be dramatically different from the original p-values. Our results show a Pearson correlation of 1 in between jackknifed p-values and the original estimates (Supplementary Figure S3). This validates that no single pathway is driving miRNA prioritization in PanomiR.

PanomiR’s implements a Gaussian estimation for miRNA-targeting p-values rather than performing bootstrapping. We showed the validity of this estimation schema by comparing the Gaussian p-values (using 1000 permutations) with bootstrapped p-values (using10K sampling). Our results show a high correlation (0.9997) between estimated and observed p-values (Supplementary Figure S4).

**Assessment pathway activity profiles**

We assessed and validated the differential pathway activity approach through two randomization schemas. In the first, we sought to measure the number of differentially activated pathways in a biologically-meaningless sample categorization by performing randomization through sample label permutation. We performed 1000 differential activation pathway experiments in the LIHC dataset with randomly assigned labels, preserving the diagnosis counts and the statistical design of the original analysis.

In the second schema, we evaluated the benefit of using annotated pathways in our differential activity analysis through generating randomized pathways. We permuted gene labels to create a pathway randomization schema that follows the structure of the annotated pathways used in this study, in terms of the number of pathways and distribution of pathways sizes. We performed 1000 gene-label permutations and performed DAP analysis using the same protocol that was performed in the LIHC dataset. We further assessed the difference between known and randomized pathways independent of conventional FDR cut-offs, via comparing the CDF of p-values using a Kolmogorov-Smirnov test. Although some randomized gene-sets were dysregulated between cases and controls at each iteration. This process showed the sensitivity of PanomiR to detect dysregulated in biologically meaningful subject classifications and low numbers of false positives in the absence of biologically meaningful signals. Pathway randomization (random gene-pathway assignments) showed an average of 693.785 differentially activated pathways at an FDR < 0.01 threshold (sd = 39.1).

**Supplementary Figures**

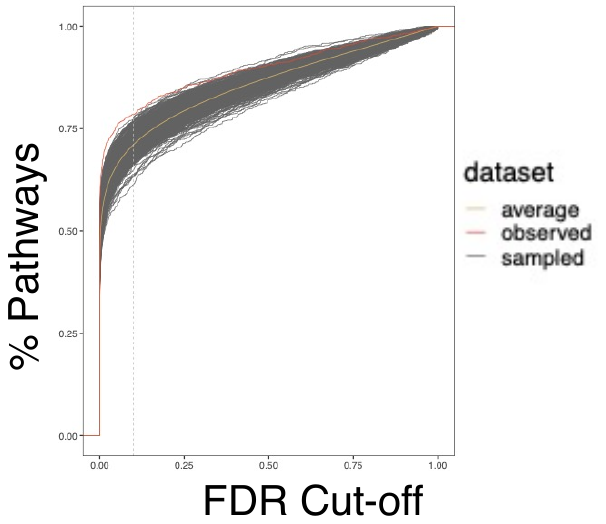

**Figure S1. Assessment of differential pathway activation analysis.** A comparison of differential pathway activity analysis in TCGA LIHC dataset using annotated and randomized pathways. The x-axis represents the FDR cut-off for determining differentially activated pathways. The y-axis represents the percentage of total pathways that are differentially activated at each cut-off. The red line is the observed number of differentially activated pathways using MSigDB Pathways (Liberzon et al. 2011). Gray lines represent datasets with randomized pathways, generated by permuting gene labels. The yellow line represents the average number of differentially activated pathways in all randomized datasets.

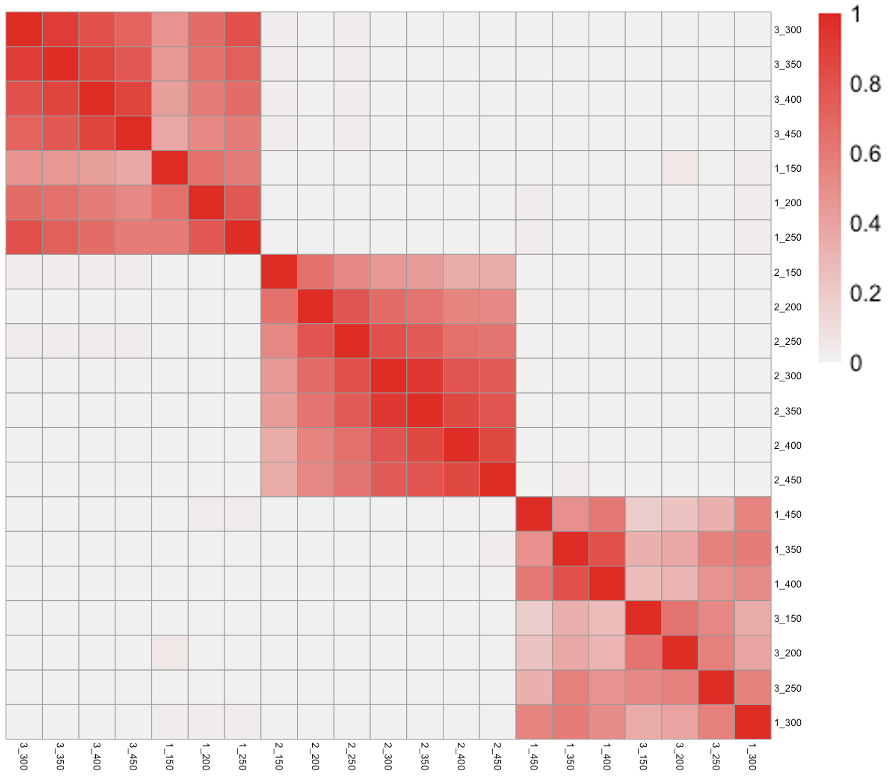

**Figure S2. The choice of number of top differentially activated pathways (DAP) does not affect cluster composition in PCxN**. Rows and columns represent differentially activated pathway clusters from PCxN using different top pathway selection criteria. At each iteration, top DAP were selected (top 150 to top 450 based on p-value; increments of 50) and mapped to PCxN. The largest 3 clusters for each iteration were identified using Louvain clustering (Cluster#_TopPathways names in rows and columns). Color intensity represents the Jaccard index of similarity between clusters. Different clusters from different iterations organize in three distinct groups, suggesting conserved groups irrespective of the number of top pathways selected.

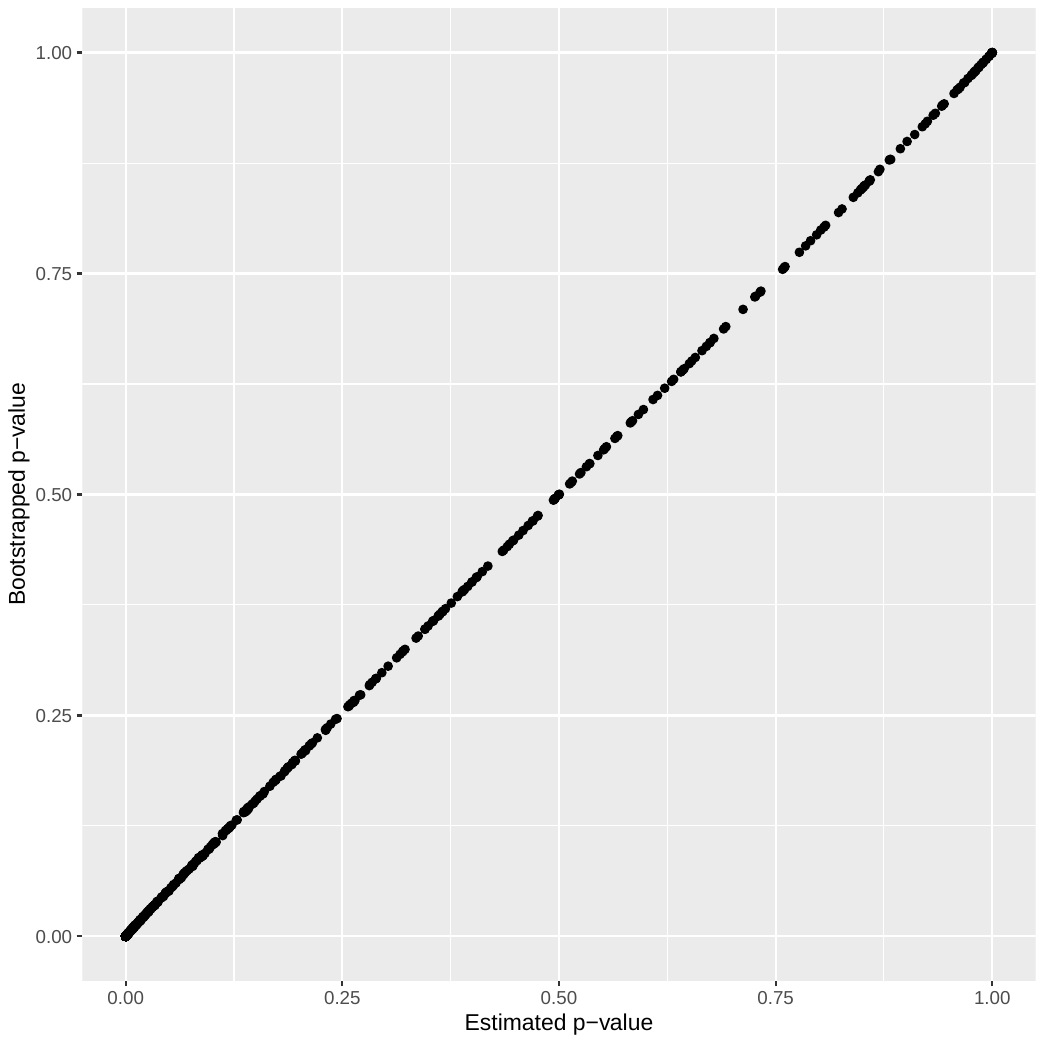

**Figure S3. Consistency of estimated PanomiR p-values with jackknifed p-values (spearman correlation = 1).** Jackknifed p-values and PanomiR p-values were generated for all miRNAs targeting the Cluster 1 of TCGA LIHC dataset using experimentally validated miRNA-mRNA interactions from the TarBase V8.0 dataset.

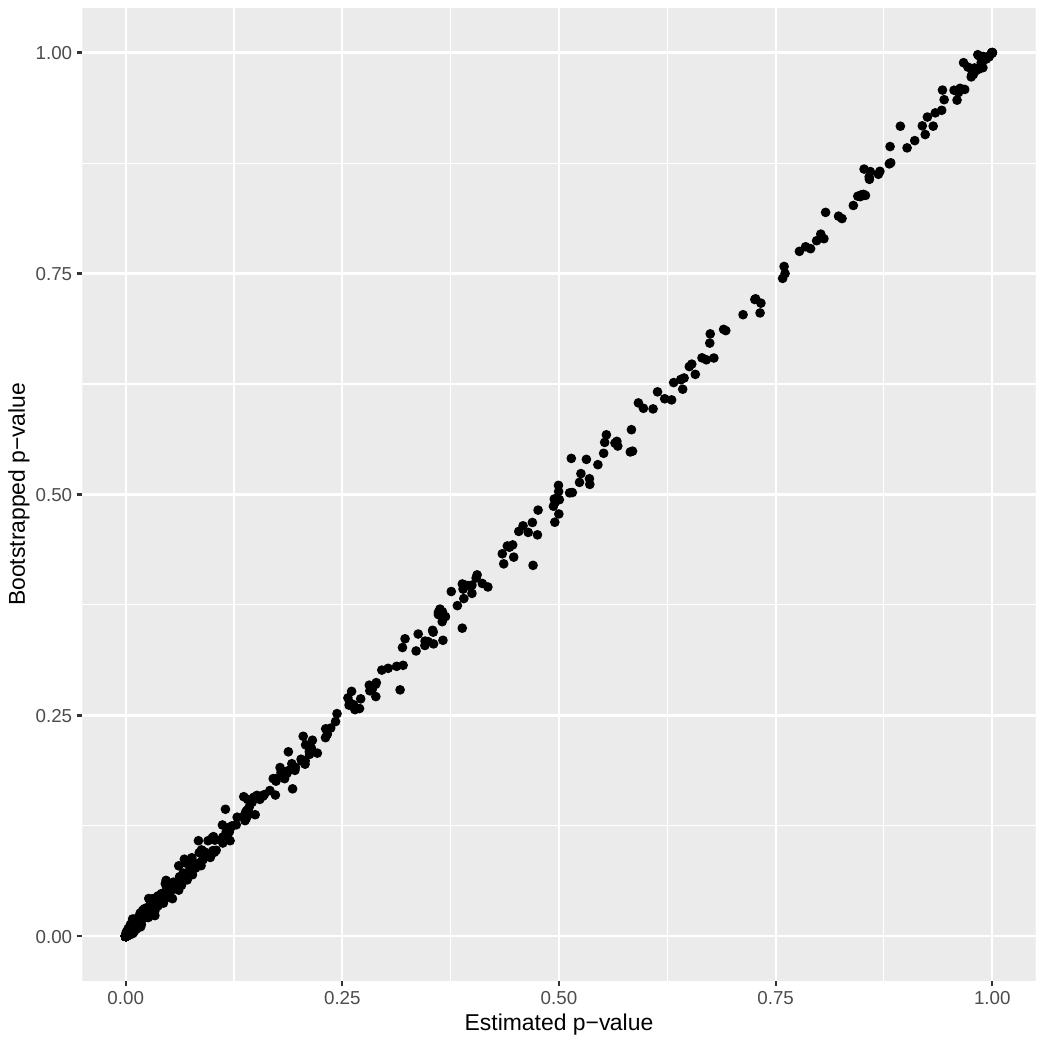

**Figure S4. Consistency of estimated PanomiR p-values with boostrapped p-values (Pearson Correlation = 0.9997).** Estimated and bootstrapped PanomiR p-values were generated for all miRNAs targeting the Cluster 1 of TCGA LIHC dataset using experimentally validated miRNA-mRNA interactions from the Tarbase V8.0 dataset (Karagkouni et al. 2018). PanomiR estimated p-values were generated from 1000 permutations and the bootstrapped were generated from 10K permutations.

**Supplementary Table Captions**

**Supplementary Table S1**. Dysregulated pathways in the TCGA liver cancer dataset. Differentially activated pathways identified by PanomiR according to p-value of differential activity between tumor (TP) and normal tissues (NT). The differential activation adjusted p-values are derived using a linear model from the limma package comparing pathway activity profiles of TP vs NT. The column Direction denotes upregulation or downregulation of pathway activity in TP vs NT.

**Supplementary Table S2. Enrichment analysis of differentially expressed genes in the TCGA liver cancer dataset.** Enrichment analysis of differentially expressed genes between TP and NT samples using MSigDB background pathways. The enrichment analysis was performed using *enricher* function from the clusterProfiler package (Yu et al. 2012).

**Table S3.** **Assessment of differential pathway activity profiling.** Differential pathway activity results were evaluated under two randomization schemas. The results are compared to readouts from the TCGA Liver Hepatocellular Carcinoma. In the pathway randomization schema, the genes in the annotated pathways were shuffled, creating randomized pathways. In the sample randomization schema, the true labels of the samples were shuffled to create artificial subject classifications. Each randomization schema included 1000 permutations.

**Table S4.** **Clusters of dysregulated pathways.** Dysregulated were mapped to PCxN and clustered using Louvain algorithm. Clusters are numbered according to their size. Clusters 1, 2, and 3 correspond to Clusters A, B, and C in Figure 4.

**Table S5.** **miRNA prioritization for the three largest clusters of differentially activated pathways, based on experimentally validated miRNA-mRNA interactions.** PanomiR prioritization of miRNAs for each identified pathway cluster, ranked by PanomiR p-value. miRNAs are prioritized based on experimentally validated miRNA:mRNA interaction from Tarbase V8.0. Enrichment analysis results are provided for comparison. The column “cluster hits” denotes the number of pathways in the cluster with significant (FDR < 0.25) enrichment in the targets of each miRNA, derived using Fisher’s Exact test.

**Table S6.** **miRNA prioritization for the three largest clusters of differentially activated pathways, based on predicted validated miRNA-mRNA interactions.** PanomiR prioritization of miRNAs for each identified pathway cluster, ranked by PanomiR p-value. (Figure 2). miRNAs are prioritized based on predicted miRNA:mRNA interaction from TargetScan V7.2. The columns “Cluster Enriched Pathways” denote the number of pathways in the cluster with significant (FDR < 0.25) enrichment in the targets of each miRNA, derived using Fisher’s Exact test.

**Supplementary Table S7. Differential expression analysis of miRNAs in the TCGA liver cancer dataset.** Differentially expressed (DE) miRNAs were identified by comparing miRNA expression in tumor tissues with normal tissues. P-values of DE miRNAs were determined using linear models provided in the limma package. Adjusted p-values were derived using FDR method.

**Supplementary Table S8. Intersection of differentially expressed miRNAs and PanomiR prioritized miRNAs using experimentally validated miRNA-mRNA interactions from Tarbase.** Differentially expressed (DE) miRNAs were identified by comparing miRNA expression in tumor tissues with normal tissues. P-values of DE miRNAs were determined using linear models provided in the *limma* package. Adjusted p-values were derived using FDR method.

**Supplementary Table S9. Intersection of differentially expressed miRNAs and PanomiR prioritized miRNAs using predicted miRNA-mRNA interactions from TargetScan.** Differentially expressed (DE) miRNAs were identified by comparing miRNA expression in tumor tissues with normal tissues. P-values of DE miRNAs were determined using linear models provided in the *limma* package. Adjusted p-values were derived using FDR method.

**Supplementary Table S10. Comparison of different parameters for selecting predicted miRNA-mRNA interactions in TargetScan database.** miRNA prioritization for the three largest clusters of differentially activated pathways, based on predicted miRNA-mRNA interactions using various parameters. The four tables represent different cut-offs for selecting miRNA-mRNA interactions. PanomiR prioritization of miRNAs for each identified pathway cluster, ranked by PanomiR p-value. (Figure 2). miRNAs are prioritized based on predicted miRNA:mRNA interaction from TargetScan V7.2. The columns “Cluster Enriched Pathways” denote the number of pathways in the cluster with significant (FDR < 0.25) enrichment in the targets of each miRNA, derived using Fisher’s Exact test.
